# A unified VCF data set from nearly 1,500 diverse maize accessions and resources to explore the genomic landscape of maize

**DOI:** 10.1101/2024.04.30.591904

**Authors:** Carson M. Andorf, Jeffrey Ross-Ibarra, Arun S. Seetharam, Matthew B. Hufford, Margaret R. Woodhouse

## Abstract

Efforts to capture and analyze maize nucleotide diversity have ranged widely in scope, but differences in reference genome version and software algorithms used in these efforts inhibit comparison. To address these continuity issues, The Maize Genetics and Genomics Database has collaborated with researchers in the maize community to offer variant data from a diverse set of 1,498 inbred lines, traditional varieties, and teosintes through a standardized variant-calling pipeline against version 5 of the B73 reference genome. The output was filtered for mapping quality, coverage, and linkage disequilibrium, and annotated based on variant effects relative to the B73 RefGen_v5 gene annotations. MaizeGDB has also updated a web tool to filter, visualize, and download genotype sets based on genomic locations and accessions of interest. MaizeGDB plans to host regular updates of these resources as additional resequencing data become available, with plans to expand to all publicly available sequence data.

## INTRODUCTION

*Zea mays* ssp. *mays* (maize, corn) has been the world’s top production grain crop for over a decade (http://faostat.fao.org/). Maize’s importance as a food, feed, fiber, and fuel product has driven its use over thousands of years, from early domestication (Hufford et al. 2012) to more directed breeding methods in the modern era (Prasanna 2012; Andorf et al. 2019).

Maize has a wealth of phenotypic, molecular, and nucleotide diversity, exceeding that of most model organisms and a great many wild and cultivated plants (Buckler *et* *al*. 2006). Assaying this diversity has proven crucial for molecular, quantitative genetic, and evolutionary studies of maize, and is central to the continued success of the maize genetics community. Efforts to capture genomic sequence diversity in maize began in 2009, when the HapMap 1 project identified 3.3 million variants in 27 diverse maize inbred lines (Gore et al. 2009). Since then, many researchers have resequenced the genomes of a diverse set of maize, providing a wealth of data for the community (Table 1).

**Table 1.**
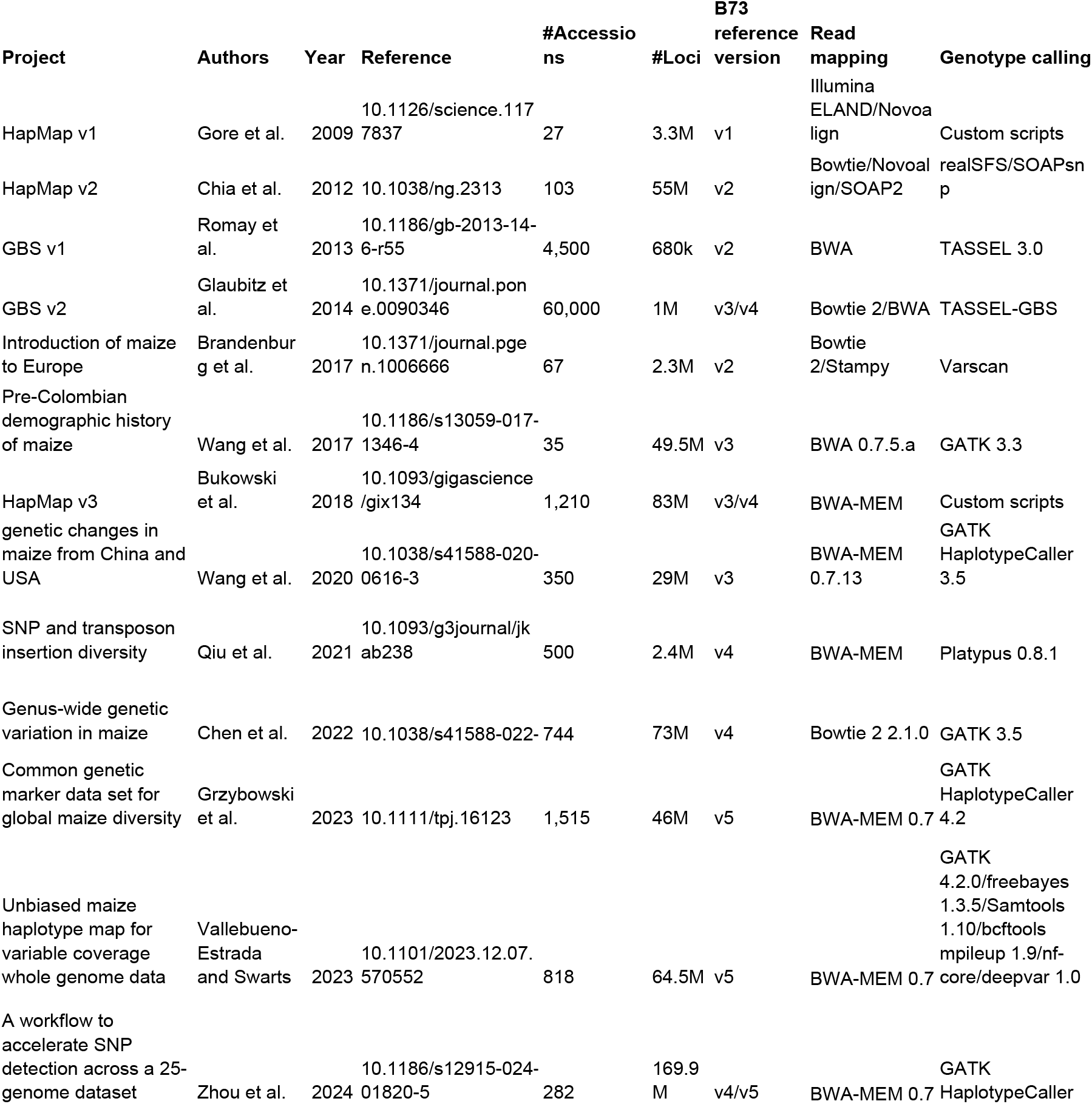
Summary of representative variant calling pipelines. Table 2. Data sets used for the WGS build.

**Table 2.** Data sets used for the WGS build.

| **Accessions** | **Project** | **Project name** | **Reference** |
| --- | --- | --- | --- |
| 539 | PRJCA009749 | WGS resequencing of 1,604 maize inbred lines | 10.1038/s41477-022-01190-2 |
| 521 | PRJNA531553 | Deep DNA resequencing of the association mapping panel | 10.1038/s41588-019-0427-6 |
| 340 | PRJNA783885 | Maize landrace whole genome resequencing | 10.1038/s41467-022-32180-9 |
| 35 | PRJEB56320 | Maize Wisconsin Diversity Panel Resequencing | 10.1111/tpj.16123 |
| 32 | PRJNA609577 | Zea mays Genome sequencing | 10.1038/s41588-020-0616-3 |
| 11 | PRJNA389800 | Whole genome sequencing of maize 282 panel | 10.1093/gigascience/gix134 |
| 8 | PRJEB14212 | European maize diversity | 10.1371/journal.pgen.1006666 |
| 7 | PRJNA399729 | Maize Haplotype Map version 3 | 10.1093/gigascience/gix134 |
| 6 | PRJNA260788 | Zea mays Genome sequencing (European maize genomes) | 10.1186/1471-2164-15-823 |
| 1 | PRJEB56265 | Resequencing of three Polish maize inbred lines | 10.1111/tpj.16123 |

In spite of the widespread application of sequencing to catalog maize diversity, the projects in Table 1 used six different mapping and eight different genotype-calling pipelines. These projects also used different versions of the maize reference genome, B73 (Hufford et al. 2021), (Jiao *et* al. 2017). Because of the divergence in pipelines and versions of the reference genome used, comparisons across these data sets are challenging, as differing versions of the reference genome result in incongruous genomic coordinates, different variant-calling pipelines can lead to conflicting variant calls, and variants may exist in one data set but not another. These continuity issues may be addressed by creating a standardized, publicly available resource, using the same variant-calling pipeline on the same reference genome.

The Maize Genetics and Genomics Database (MaizeGDB – https://www.maizegdb.org) (Woodhouse et al. 2021) is the maize community database providing data curation and informatics resources to support genetics, genomics, and breeding research for maize scientists. Here, MaizeGDB has collaborated with researchers in the maize community to offer variant data through a standardized genotype-calling pipeline that uses BWA-MEM (Li 2013) and Sentieon’s Haplotyper and GVCFtyper (Freed et al. 2017) against version 5 of the B73 reference genome. MaizeGDB offers the entire data set as a download, and we describe updates to the SNPversity application (Schott et al. 2018) that allow users to filter, visualize, and download variants based on their selected filters and parameters. MaizeGDB has also integrated these data into the tool PanEffect (Andorf et al. 2023), which visualizes and predicts the variant effects of missense mutations in maize accessions relative to the reference B73 v5 genome.

## METHODS

### Gencove pipeline

The MaizeGDB WGS build was performed by Gencove https://gencove.com/. Reference assembly Zm-B73-REFERENCE-NAM-5.0 (NCBI RefSeq Assembly GCF_902167145.1) was cURLed from https://download.maizegdb.org/Zm-B73-REFERENCE-NAM-5.0/Zm-B73-REFERENCE-NAM-5.0.fa.gz. BWA indices of fastq files https://bio-bwa.sourceforge.net/ were created using Sentieon Driver’s (version 202010.01) (Freed et al. 2017) optimized implementation of BWA-MEM bwa index -a bwtsw. An index Zm-B73-REFERENCE-NAM-5.0.fa of was created using samtools (version 1.15.1) faidx http://samtools.sourceforge.net/ (Danecek et al. 2021) and a sequence dictionary was created using Picard’s (version 3.0.0) CreateSequenceDictionary http://picard.sourceforge.net/. High-depth paired-end fastq files (n=1498) from the accessions described below were downloaded from various public repositories (Table 2, Supplemental Table 1). The paired-end sequences were aligned to the reference assembly using Sentieon Driver’s (Freed et al. 2017) BWA-MEM. Variant calling was performed using Sentieon’s Haplotyper with gVCF output per sample. Joint variant calling was performed using Sentieon’s GVCFtyper, and a maximum of 12 alternate alleles are reported for each site. The single-output VCF file contains only variant sites.

### Post-processing pipeline

The initial joint VCF was first filtered with a custom Python script to retain reads with mapping quality over 30 and coverage over 50% and remove multiallelic variants. Next, variant effects were annotated to the B73 v5 gene models using SNPEff (version 5.2). This preprocessing yielded a set named the “MaizeGDB 2024 High-Coverage” data set, comprising 228,679,451 variants, with roughly 12.5 million variants located within genic regions of the genome. A subsequent filtering phase produced the “MaizeGDB 2024 High-Quality” data set. Plink (version 1.9) was used to calculate pairwise linkage disequilibrium (R^2^). Only variants with a maximum R^2^ of 0.5 or greater, in relation to at least one other variant within a distance of 400-5000 bp, were retained. This refinement resulted in a collection of 75,518,390 variants, approximately 6.3 million of which are located in genic regions of the genome. Both the high-coverage and high-quality data sets are accessible in VCF format. Finally, a custom Python script using the h5py library converted the VCF files to HDF5 format for easier storage and retrieval. Filtering types and statistics can be found in Table 3. The scripts and commands used in post-processing are available on the GitHub page (https://github.com/Maize-Genetics-and-Genomics-Database/SNPversity2.0).

### MaizeGDB SNPversity 2.0 tool

SNPversity 2.0 (https://wgs.maizegdb.org/) (https://github.com/Maize-Genetics-and-Genomics-Database/SNPversity2.0), a complete rewrite of the original SNPversity tool released by MaizeGDB in 2018 (Schott et al. 2018), represents a significant upgrade designed to enhance the exploration of genetic variation within local genomic regions. Central to this new version is its reliance on an HDF5 database back-end, optimized to efficiently manage variant annotations. SNPversity 2.0 uses a Python-based data exchange layer and a JavaScript-driven interface.

This combination facilitates rapid access to variant data, enabling the real-time presentation of single nucleotide polymorphisms (SNPs) and insertions/deletions (INDELs) in a highly accessible and interactive manner.

## RESULTS AND DISCUSSION

The MaizeGDB WGS build is composed of a diverse set of inbred lines, landraces, and teosintes from 1,498 resequenced accessions mapped to version 5 of the reference B73 genome. The accessions are from ten bioprojects (Table 2, Supplemental Table 1), and include data from the maize 282 panel, the European diversity panel, HapMap 3, a large Chinese association mapping panel, elite inbreds from China, and traditional varieties from Mexico. The filtered WGS data can be analyzed through MaizeGDB’s SNPversity 2.0 variant processing tool (https://wgs.maizegdb.org/) (Figure 1). SNPversity is an open-source, web-based variant visualization tool originally released by MaizeGDB in 2018 (10.1093/database/bay037). Users can specify genomic intervals and select maize accessions of interest, with the system producing outputs in the form of Variant Call Format (VCF) files, detailed data tables, and neighbor-joining tree views. These outputs display alleles corresponding to the user’s queries with improved data interpretation via color-coded tables showing each SNP’s allelic status and mutational state. New to SNPversity 2.0, the platform now offers valuable metadata such as variant effect annotations, mapping quality and coverage scores, linkage disequilibrium measures, and interoperability with the MaizeGDB genome browsers and PanEffect tool.

**Figure 1.**
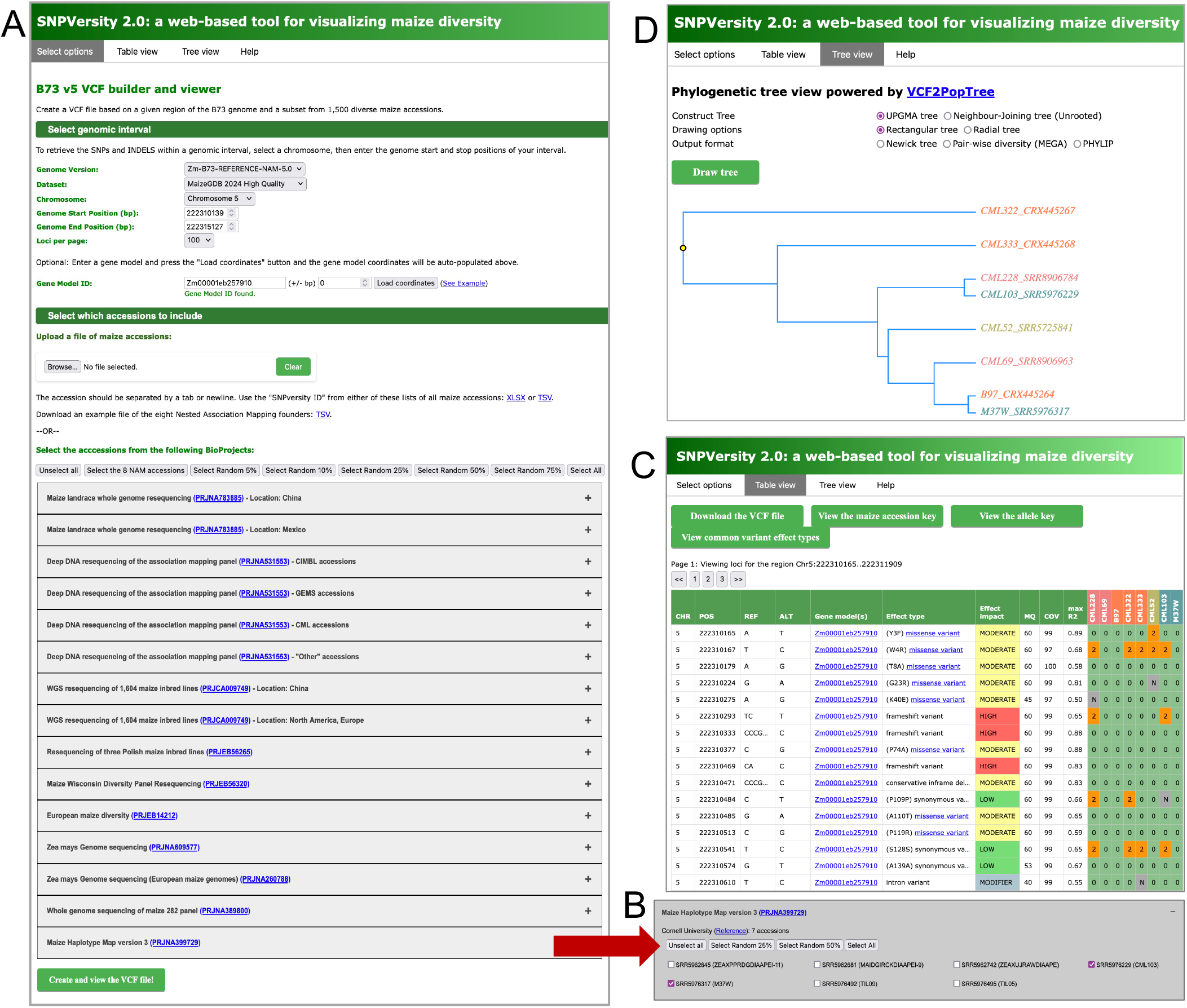
Overview of SNPversity 2.0. A) The “Select options” tab. B) An example of one of the projects, opened to view the accessions within and the option selections. C) The “Table view” output tab featuring eight NAM founder accessions for gene model Zm00001eb257910. D) The “Tree view” output tab of eight NAM founder selections for gene model Zm00001eb257910.

SNPversity 2.0 data are organized into three tabs (Figure 1): “Select options” (Figure 1A), “Table view”(Figure 1B), and “Tree view” (Figure 1D). The “Help” tab includes detailed instructions on how to use the SNPversity 2.0 tool.

### SNPversity 2.0 “Select options” tab

The “Select options” tab is the B73 v5 VCF builder and viewer. The top subsection, “Select genomic interval”, retrieves the variants within a genomic interval of interest, while also selecting how many loci should appear per page. A user can select which MaizeGDB filtered data set to query: the “MaizeGDB 2024 High-Quality” data set, or the “MaizeGDB 2024 High-Coverage” data set. A user may also enter a B73v5 gene model ID to restrict the search to that gene model’s genomic interval: the user enters the gene model ID in the “Gene Model ID” row, then clicks the “Load coordinates” button. Users have the option to add or subtract base-pair coordinates from the gene model start-stop coordinates in the “(+/-bp)” window. There are various ways to select or filter accessions in the “Select which accessions to include” subsection, including uploading a file with a list of accessions of interest; selecting accessions by Bioproject; by randomly generated percentages of all Bioprojects; or by randomly generated percentages of a specific Bioproject. Users can also select the eight available Nested Association Mapping (NAM) founder lines (McMullen et al. 2009) for processing. The “Create and view the VCF file” button at the bottom of the page generates the output. Outputs for a user’s selection will appear in the “Table view” tab at the top.

### “Table view” tab

The “Table view” tab contains the outputs of the variant data sets and parameters selected in the “Select options” tab (Figure 2). Figure 2A shows the variant output of the genomic region of gene model Zm00001eb257910 and the alleles of the eight accessions associated with the maize Nested Association Mapping (NAM) founder population. Outputs are organized by 1) chromosomal coordinates; 2) reference and alternate allele; 3) the gene model located within the coordinates; 4) the effect type, with missense changes indicated in a single-letter amino acid substitution notation (e.g. for the example Y3F, Y=the amino acid for the reference genome, 3=the amino acid position, and F=the amino acid substitution in the alternative genome); 5) the predicted effect impact; 6) mapping quality (MQ); 7) coverage (COV); and 8) max R^2^ within 400-5000bp; followed by the variant scores across the selected accessions (see Methods for how these factors were calculated). Variant scores are labeled either 0 (reference allele), 1 (heterozygous alternate allele), 2 (homozygous alternate allele), or N (no data). The button “Visualize common variant effect types” at the top right describes the definitions of the variant effects (such as frameshift variant) and effect impacts (low, moderate, or high). Accessions are organized and color-coded by Bioproject. The button “View the maize accession key” at the top center describes which project is associated with each color. Above the table, users can select which page of data to navigate to, if there are multiple pages. The data in the table can be downloaded as a vcf file by clicking on the “Download the vcf file” button at the top left.

**Figure 2.**
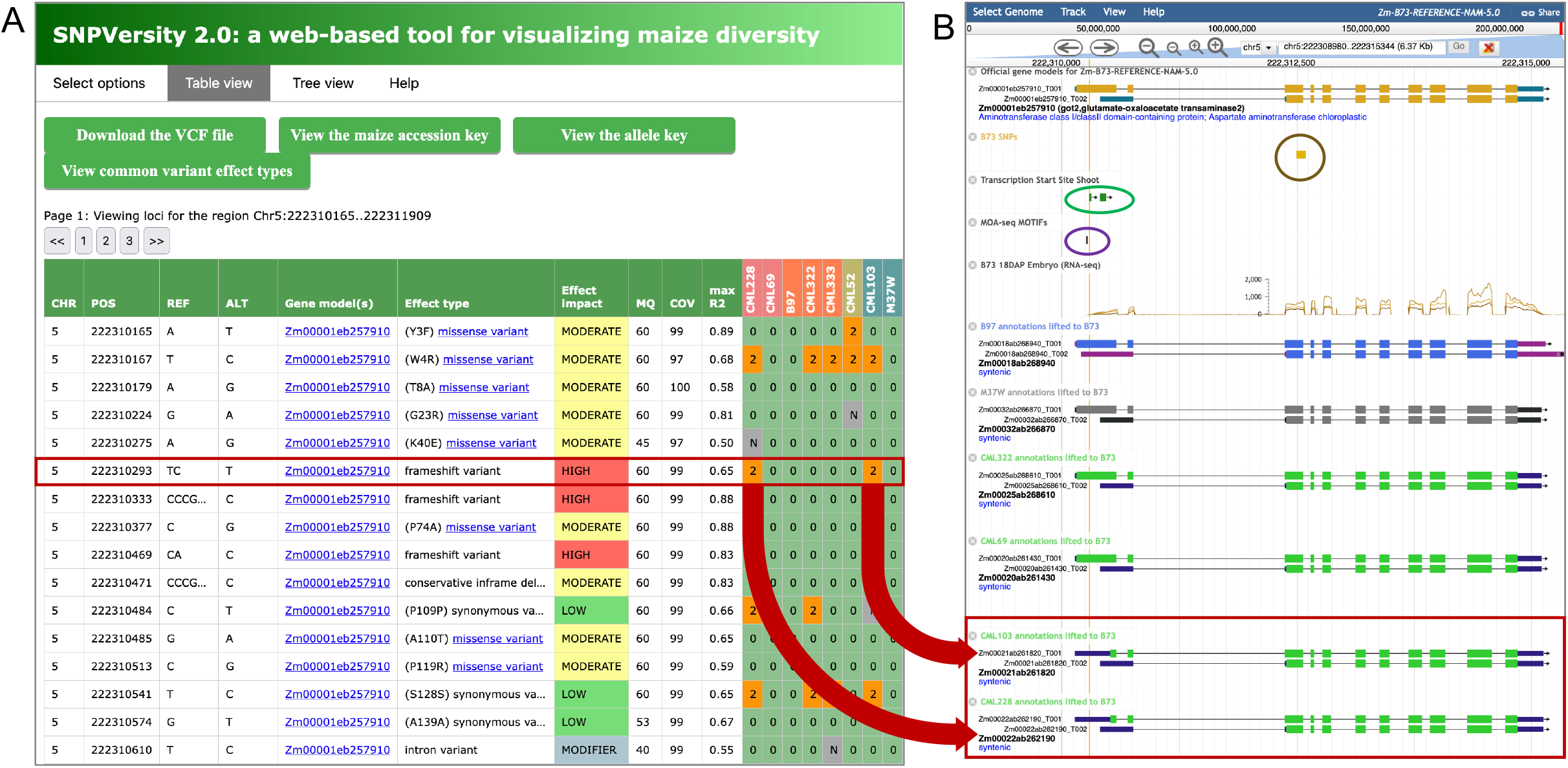
Example data in the SNPversity 2.0 “Table view” tab. A) Eight NAM founder accessions for the gene model Zm00001eb257910. B) JBrowse instance of the variant highlighted in red from A. Featured are the reference B73 gene model Zm00001eb257910 at the top, followed by the kernel GWAS locus, circled, the shoot transcription start site, circled, the MOA-seq motif, the 18DAP RNA-seq track, and the lifted over gene model annotations to genomes corresponding to the NAM founder accessions from A. The red arrows from A of CML228 and CML103 that have the TC->T variant point to the lifted over gene models from their respective genomes. The TC->T variant from A is represented by the yellow vertical line. The frameshift variant corresponds to a truncated 5’ exon for CML228 and CML103 relative to the other gene models.

Linking the variant data to the JBrowse instances in MaizeGDB enables users to perform robust functional analyses of their loci of interest. Figure 2A highlights a variant at position 222,310,293 on chr5 that results in a TC->T frameshift variant with a predicted high effect impact. Accessions CML228 and CML103 have the alternate TC->T variant. Clicking on the hyperlink for the gene model ID in the Gene Model(s) column will take the user to the exact location of the reference allele in the B73 v5 JBrowse genome browser, shown in Figure 2B. This example shows that this variant, shown by the vertical yellow line, is located in the 5’ exon of Zm00001eb257910.

This gene model is the glutamate-oxaloacetate transaminase2 gene *got2* (https://maizegdb.org/gene_center/gene/12270). The JBrowse instance of *got2* in Figure 2B shows that this gene has been identified as a kernel weight GWAS candidate (Wallace *et* *al*. 2014). The location of the frameshift variant is associated with a CAGE-identified shoot transcription start site (Mejía-Guerra et al. 2015), and it is adjacent to a maize ear MOA-seq (MNase-defined cistrome-Occupancy Analysis) motif that identifies a putative transcription binding motif (Savadel et al. 2021). An RNA-seq track of 18 days after pollination embryo RNA-seq reads initiates at the frameshift variant location. Below are gene model annotations lifted over to B73 from some of the accessions shown in Figure 2A (Hufford et al. 2021; Woodhouse et al. 2021). The accessions (CML228 and CML103) that contain the variant, have a truncated 5’ exon relative to the Zm00001eb257910_T001 transcript. Together, these data suggest that the variant TC->T for CML228 and CML103 resulted in a truncated gene model for both genomes.

The missense variant data from the MaizeGDB build have also been integrated into MaizeGDB’s PanEffect tool (Andorf et al. 2023) (Figure 3). PanEffect visualizes and predicts the protein sequence missense variant effects in maize accessions relative to the reference B73 v5 genome. To access all predicted variants in the PanEffect tool from SNPversity 2.0, users can select the “MaizeGDB 2024 High-Coverage” data set from the “Select options” tab, Figure 3A. Figure 3B shows the output of the High-Coverage data set for gene model Zm00001eb257910. Highlighted in red is a missense variant from P->L at codon 296 (P296L). Clicking on the “missense variant” link under the “Effect type” column will navigate the user to PanEffect (3C).

**Figure 3.**
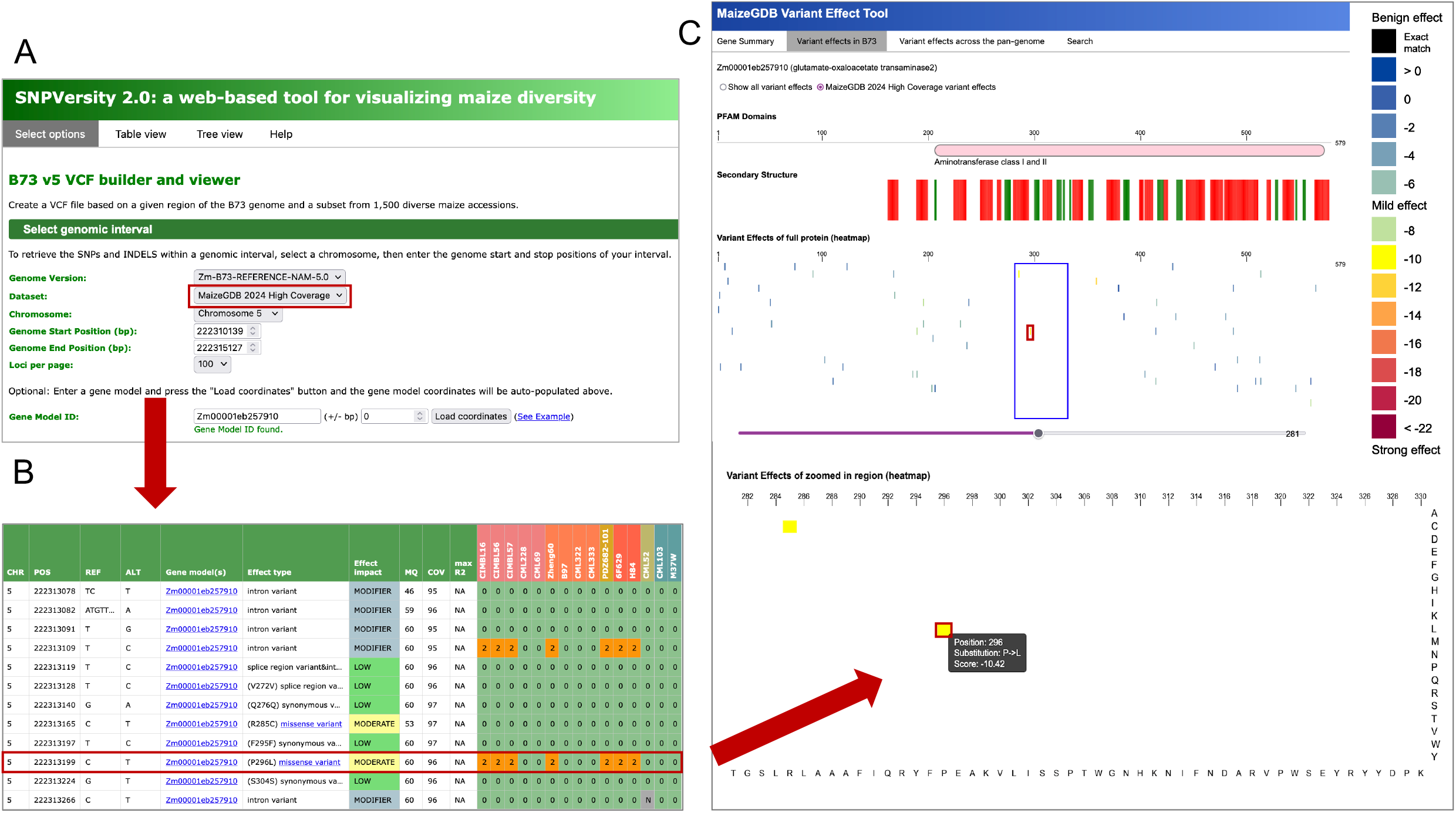
Connecting SNPversity 2.0 to the MaizeGDB PanEffect tool. A) select the “High-Coverage” data set from the SNPversity 2.0 “Selection options” tab. B) High-coverage output for gene model Zm00001eb257910. Click on the “missense variant” link in the “Effect type” column to go to the predicted codon missense variant effect prediction in PanEffect. C) The PanEffect visualization platform. By selecting the “Variant effects in B73” tab and the “MaizeGDB 2024 High Coverage variant effects” radio button, the missense variants from the High-Coverage WGS data are shown, with color-coded effect predictions for each missense variant, from blue (benign effect) to red (strong effect). Circled is the P->L missense variant at codon 296 (P296L) for Zm00001eb257910 from SNPversity 2.0 shown in Figure 3B.

By selecting the “Variant effects in B73” tab and the “MaizeGDB 2024 High Coverage variant effects” radio button, the missense variants from the High-Coverage WGS data are shown, with color-coded effect predictions for each missense variant, from blue (benign effect) to red (strong, likely negative, effect). Circled is the P->L at codon 296 (P296L) from SNPversity 2.0 shown in Figure 3B; colored yellow, it is predicted to have a mild to moderate effect. In this example, accessions outside of the eight NAM accessions were selected in SNPversity 2.0 to show accessions that have the variant, since the NAM accessions did not.

### “Tree view” tab

The “Tree view” tab (Figure 1D) allows users to construct a UPGMA tree or a neighbor-joining tree (Unrooted) from the “Table view” output via VCF2PopTree (Subramanian et al. 2019). Users can draw a rectangular or radial tree, or output a Newick tree, Pairwise diversity (MEGA), or PHYLIP format. Accessions in the tree visualization for the SNPversity 2.0 “Tree view” tab are color-coded by project based on the schemes shown in the “Table view” tab.

## SUMMARY

A standardized, publicly available resource using a single pipeline to map and call variants across a diverse range of maize accessions will help maize researchers to perform comparative genomics experiments without facing the continuity issues that exist when attempting to compare data sets from disparate pipelines. MaizeGDB offers this entire data set as downloads and through SNPversity 2.0 to filter, visualize, and download variants based on user-selected filters and parameters and plans to update the data set on a regular basis as new accessions become available.

## WEB RESOURCES

The scripts used to process the WGS data and descriptions of the filtering process can be found at https://github.com/Maize-Genetics-and-Genomics-Database/SNPversity2.0. The “Help” tab for SNPversity 2.0 includes an in-depth description of how to use the tool.

## DATA AVAILABILITY STATEMENT

The WGS build used accessions from Bioprojects PRJCA009749, PRJEB14212, PRJEB56265, PRJEB56320, PRJNA260788, PRJNA389800, PRJNA399729, PRJNA531553, PRJNA609577, PRJNA783885. All data are publicly available at GenBank https://www.ncbi.nlm.nih.gov/nucleotide/, except for PRJCA009749, which is available at the Genome Sequence Archive https://ngdc.cncb.ac.cn/gsa/. Individual accessions are listed in Supplemental Table 1. The filtered WGS data can be downloaded from https://ars-usda.app.box.com/v/maizegdb-public/folder/255390517505. The full, unfiltered WGS data is available upon request.

## ACKNOWLEDGEMENTS

We acknowledge the smallholder farmers and indigenous people whose work and love for their traditions and identity keep maize diversity alive. This research was supported by the US. Department of Agriculture, Agricultural Research Service, Project Number [5030-21000-072-00-D] through the Corn Insects and Crop Genetics Research Unit in Ames, Iowa. The work for this project was also conducted through USDA-ARS non-assistance cooperative agreement Accession 442724, Iowa State University. This research used resources provided by the SCINet project and the AI Center of Excellence of the USDA Agricultural Research Service, ARS project numbers 0201-88888-003-000D and 0201-88888-002-000D. J.R.-I. acknowledges support from the USDA Hatch project CA-D-PLS-2066-H. Mention of trade names or commercial products in this publication is solely for the purpose of providing specific information and does not imply recommendation or endorsement by the U.S. Department of Agriculture. USDA is an equal opportunity provider and Employer. Finally, we would also like to acknowledge Felix Andrews for a rousing discussion of whether we should name the data set HapMap π or HapMap IV.

